# Prefrontal cortex regulates amygdala response to threat in trait anxiety

**DOI:** 10.1101/215699

**Authors:** Maria. Ironside, Michael. Browning, Tahereh L. Ansari, Christopher J. Harvey, Mama N. Sekyi-Djan, Sonia J. Bishop, Catherine J. Harmer, Jacinta O’Shea

## Abstract

**Background:** Highly co-morbid mood and anxiety disorders are associated with aberrant fronto-limbic signalling during emotional processing. Animal models suggest that hypoactive prefrontal cortex weakens top-down control of limbic structures, causing heightened limbic and behavioural reactivity to negative information. Here we tested for this causal mechanism in human trait anxiety. We reasoned that if dorsolateral prefrontal cortex controls amygdala response to affective information, then stimulation of that brain region should reduce the hyperactive amygdala threat responsivity seen in trait anxiety.

**Methods:** Using a within-subjects design, sixteen high-trait anxious females received active and sham transcranial direct current stimulation (tDCS) of the dorsolateral prefrontal cortex, in counterbalanced order, with sessions timed to be at least one month apart. Each session was followed immediately by a functional magnetic resonance imaging (fMRI) scan during which participants performed an attentional task with threat-related distractors.

**Results:** As predicted, compared to sham stimulation, active prefrontal cortex stimulation reduced amygdala threat reactivity and simultaneously increased activity in cortical regions associated with attentional control and improved task accuracy.

**Conclusions:** These results demonstrate a causal role for impoverished frontal regulation of amygdaloid function in attentional capture by threat in trait anxiety. The finding that prefrontal stimulation reduces amygdala threat reactivity acutely indicates a neurocognitive mechanism that could contribute to tDCS treatment effects in affective disorders.

## Introduction

The difficulty in treating highly co-morbid mood and anxiety disorders(1) has increased clinical interest in potential alternative treatments, such as non-invasive brain stimulation techniques, including transcranial direct current stimulation (tDCS) of the dorsolateral prefrontal cortex (DLPFC). A recent meta-analysis(2) of individual patient data from 289 tDCS-treated patients showed that, compared with placebo-controlled clinical trials of antidepressant drugs, sham-controlled clinical trials of prefrontal tDCS have a similar number needed to treat (7 for response, 9 for remission). These results highlight the potential value of prefrontal tDCS in the treatment of affective disorders.

However, as with many antidepressant treatments, the mechanism of action is unclear. The DLPFC is implicated in animal studies, which provide compelling evidence of the importance of the medial prefrontal cortex in regulating responses to threat via direct inhibition of the amygdala complex. The prefrontal cortex has been shown to inhibit aversive associations established in fear conditioning, with prefrontal lesions impeding(3) and prefrontal electrical stimulation enhancing(4) the extinction of a conditioned response in mice and rats. Furthermore, electrical pre-stimulation of the prefrontal cortex in rats and cats(5) specifically blocks or reduces amygdala responses. The results from these preclinical investigations provide the foundation for theoretical models of emotional dysfunction in disorders of anxiety and depression, where deficient prefrontal control is believed to result in over-activity within areas responsible for assigning salience and attention to biological stimuli, such as the amygdala.

It is well established that the amygdala is a critical component of the neural circuitry underlying fear processing(6). Consistent with this, human neuroimaging studies have confirmed hyperactive amygdala and/or hypoactive prefrontal cortex activity in patients with anxiety disorders(7) and major depressive disorder(8, 9), indicating an imbalance of activity within this cortico-limbic circuit in human affective disorders. Furthermore, there is evidence that treatment with antidepressant drugs(10) or cognitive behavioural therapy(11) can reduce amygdala hyperactivity. Antidepressant treatment with deep brain stimulation of the subgenual cingulate increases DLPFC activation(12), and recent functional connectivity data suggest that invasive and non-invasive forms of brain stimulation may recruit overlapping functional networks (13). However, there is no direct causal evidence that the prefrontal-amygdala circuit functions in humans as reported in preclinical animal models, i.e. that prefrontal cortical control regions inhibit amygdala responses to threat. Here we combined tDCS with functional magnetic resonance imaging (fMRI) to perform a causal test of this hypothesis in a high trait anxious sample.

TDCS can be used to tonically increase or decrease cortical excitability using weak electrical currents(14). Induced changes in tissue excitability can persist over minutes to hours after stimulation (depending on stimulation current, duration and number of sessions), effects that are N-methyl-D-aspartate (NMDA) receptor-dependant, and presumed to reflect changes in synaptic efficacy and plasticity(15, 16). Initially used as a tool to induce changes in motor evoked potentials, tDCS has more recently been used to modulate cognition, such as attentional control(17) and working memory(18). Neuroimaging indicates that prefrontal tDCS alters functional activation and connectivity in brain regions that support cognitive function, including regions distal from the stimulating electrodes(19, 20)(21). Thus, the mechanism of action of prefrontal tDCS in the treatment of depression may arise through the induction of plasticity in distributed cortical-striatal/limbic circuits, a network hypothesis that can only be assessed through combined neuro-stimulation/imaging research. Using stimulation to change the electrical state of cortical tissue, it becomes possible to test causal hypotheses about functional interactions between cortex and connected subcortical structures(22), such as test the causal influence of pre-frontal cortex on regulating the amygdala response to threat.

We have shown behaviourally that bilateral prefrontal tDCS reduces vigilance to threat in an attentional task validated to predict the clinical response to anxiolytic drug treatment(23). We hypothesize that this effect arises because prefrontal stimulation increases cortical activity driving top-down attentional control of connected limbic structures, thus increasing regulation of the amygdala threat response. To test this mechanistic hypothesis, here we assessed the effect of prefrontal tDCS in a high trait anxious group on neural threat reactivity measured with fMRI, during a well-validated attentional control paradigm (perceptual load-face paradigm with neutral and fearful face distractor stimuli) that is sensitive to anxiety-related differences in attentional function. Previous work with this task has shown that high anxious individuals exhibit a hypoactive frontal cortex response and hyperactive amygdala response to fearful face distractors(24). Crucially, this was apparent only under conditions of ’low attentional load’. That is, when the task is undemanding and does not fully occupy attentional resources, high anxious individuals exhibit impoverished frontal recruitment and increased amygdala responsivity to threat-related distractors. However, the causal relationship between this concurrent frontal hypo-responsivity and amygdala hyper-reactivity has not been determined. Here, we sought to address this using tDCS of the prefrontal cortex as a causal intervention in a high trait anxious group. We predicted that prefrontal tDCS would modulate this pattern of activation and behaviour.

Specifically, we predicted that, under conditions of low attentional load with fearful distractors, tDCS would have three directional effects: 1) *increase* cortical activation; 2) *decrease* amygdala activation; and 3) *improve* task accuracy. We stimulated bilateral prefrontal cortex and then assessed the causal impact on brain activity and behaviour. As predicted, under conditions of low attentional load, tDCS both increased activity in cortical regions associated with attentional control and reduced amygdala activity, specifically on fearful distractor trials. tDCS also tended to improve accuracy on fearful distractor trials.

## Methods and Materials

### Participants

The study was approved by the Central University Research Ethics Committee (University of Oxford) and performed in compliance with their approved protocols. Owing to higher prevalence of trait anxiety in the female population, a sample of eighteen female participants (all right handed, aged 18-45 years, mean 23 years) was recruited from the community. Participants were pre-screened with an online version of the Spielberger State-Trait Anxiety Inventory (STAI(25)) and those who scored >45 on the trait anxiety questionnaire (STAI-T) (a conservative cut-off, as scores above 39-40 reflect clinically significant symptoms(26)) were invited to a screening session at the Warneford Hospital, Oxford, where they completed the Structured Clinical Interview for DSM-IV disorders(27). Participants STAI-T scores ranged from 45 to 63 (mean = 53, standard deviation [SD] = 5). Written informed consent was obtained from all participants. Individuals with current depressive episode, current or past neurological disease, or family history of bipolar disorder were excluded, as were individuals on medication for anxiety or depression, or with any contraindications to MRI or tDCS. Participants who successfully met full screening requirements were invited to take part in two tDCS/MRI scanning sessions at the John Radcliffe Hospital, Oxford. Participants were compensated for their time at a rate of £10 per hour. Formal sample size calculation was precluded, because no prior study had determined the effect of tDCS on brain activity in a high anxious sample. Hence, we estimated the likely effect size of tDCS, and the likely minimum sample size, informed by two prior related studies. Our previous work(23) showed that prefrontal tDCS reduced behavioural threat vigilance in healthy volunteers, with an effect size of Cohen’s d_s_ = 0.87(28). Another previous work, using fMRI and the identical task paradigm to that used here, reported higher amygdala and lower prefrontal cortex activation in a high versus low anxious sample(24), with an effect size of Cohen’s d_s_ =0.99. Informed by these data, *a priori* sample size calculation for the current repeated measures design (29) yielded n=8 as the minimum sample size required to detect a reduction in amygdala fMRI signal of this magnitude (difference between two dependent means: matched pairs, one tailed, alpha=.05, dz = 0.99, power=.8), and n=10 to detect a reduction in attention to threat behaviour of this magnitude (difference between two dependent means: matched pairs, one tailed, alpha=.05, dz = 0.87, power=.8). For the present study we recruited sixteen participants, Data for two participants were partially lost due to server problems. These two participants were replaced to bring the total N recruited to eighteen and the dataset analysed to sixteen.

### Design

This study used a within-subjects double-blind design with 16 participants each attending two separate tDCS/fMRI sessions, randomised to stimulation order (real/sham tDCS followed by sham/real tDCS one month later, counterbalanced). A randomisation list was prepared by a colleague (Department of Psychiatry, University of Oxford) separate from the study and kept in a locked cabinet. Based on this randomisation list the experimenter was given a code to enter into the tDCS device which determined whether real or sham stimulation was delivered. Thus, the experimenter was blind to the stimulation order. On the day of the study participants first filled out mood questionnaires before being introduced to the scanner environment and undergoing a structural scan, during which they practiced a training version of the attentional control task. Then participants vacated the scanner and received tDCS in a separate room while they sat at rest. This allowed the participants to practice the task and become comfortable in the scanner environment to minimise time spent entering the scanner after tDCS. After the stimulation ended the participants were reintroduced into the scanner (mean time from tDCS offset to task onset ~7 minutes) and carried out the attentional control task while fMRI data were acquired.

### Attentional load paradigm

Visual stimuli were back projected onto a translucent screen positioned behind the bore of the magnet, visible via an angled mirror placed above the participant’s head. The attentional load paradigm (and following task description) was adapted from Bishop(24) and others. In each trial, a string of 6 letters superimposed on a task-irrelevant non-familiar face was presented for 200 ms (see Fig.1). In the present study the face stimuli comprised four different individuals with fearful and neutral expressions taken from the Pictures of Facial Affect(30) and cropped to remove extraneous background information. The neutral faces were morphed using computer graphics to have a neutral: happy expression mix of 30:70%, because wholly neutral faces have previously been found to be aversive(31). The experiment was performed using Presentation^®^ software (Version 14.0, Neurobehavioral Systems, Inc., Berkeley, CA).

**Figure 1.**
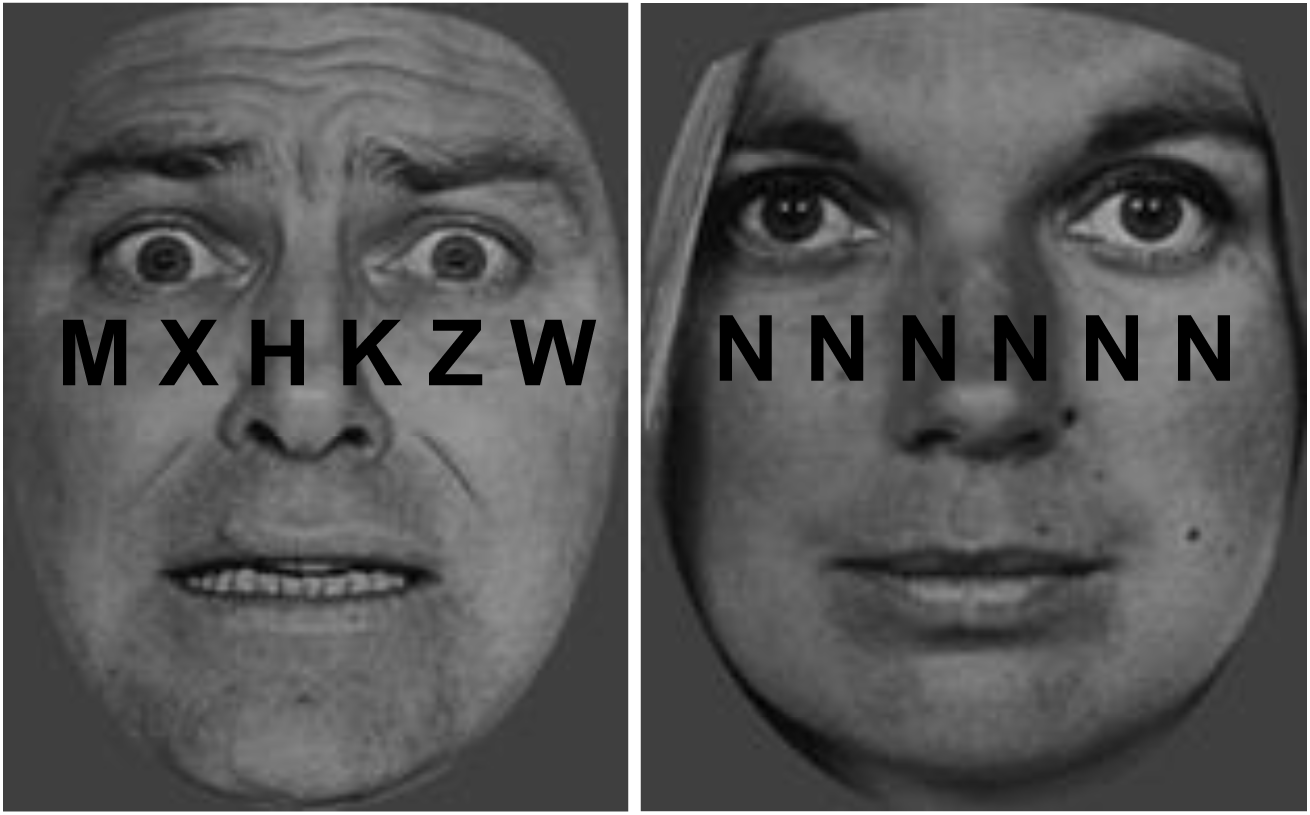
Example stimuli in the attentional load paradigm. On each trial, a string of 6 letters was superimposed on a task-irrelevant face distractor presented in the centre of the screen. Face stimuli reproduced with permission from Ekman and Friesen (1975). Left hand side – fearful distractor, high load; right hand side - neutral distractor, low load. Participants had to indicate whether the letter string contained an “X” or an “N”. A target was present on every trial.

The task was to decide whether the letter string contained an “X” or an “N”. In half the blocks – the *“high attentional load”* condition – the string comprised a single target letter (N or X) and 5 non-target letters (H, K, M, W, Z) arranged in random order. In the other half of blocks—the ‘‘*low attentional load*" condition—the letter string comprised 6 Xs or 6 Ns, reducing attentional search requirements. This manipulation of attentional load is identical to the one used in Bishop and others(24), Jenkins and others(32) and conforms to Lavie’s(33) description of heightening cognitive effort by: 1) increasing the number of different identity items that need to be perceived, or 2) making perceptual identification more demanding on attention. The rationale for these load conditions is that when the task is undemanding, greater distractibility puts higher demands on attentional control.

A mixed block/event-related design was used — the level of attentional load (high or low) was varied across blocks, while the expression of the task-irrelevant face distractors (fearful or neutral) was varied across trials. These 2 factors (attentional load × distractor emotion) resulted in 4 conditions: high load/fearful distractors; high load/neutral distractors; low load/fearful distractors; low load/neutral distractors. The key hypothesis-driven condition of interest was: low load/fearful distractors. Previous work has shown that amygdala response to threat is observed only in the low load condition in this task(24). Therefore, by examining the effect of tDCS on brain regions selectively activated by this key hypothesis-driven contrast (fearful versus neutral face distractors under conditions of low load) it was possible to test the hypothesis that tDCS reduces vigilance to threat in trait anxiety by altering fronto-limbic activity; specifically, by reducing amygdala response to fearful distractors.

There were 3 imaging acquisition runs, each comprising 12 blocks of 4 trials. There was a 2 s interval between blocks. Within blocks, the inter-stimulus interval was randomly jittered using an exponential function with a mean of 4.5 s and a minimum of 3 s.

### Transcranial Direct Current Stimulation (tDCS)

Stimulation was delivered using a battery powered device (DC Stimulator Plus, Neuroconn, Germany(34)). The rubber electrodes (5cm × 5cm) were placed in saline soaked sponges and affixed to the scalp with a rubber band. We used a bipolar-balanced electrode montage which positioned the anode (positive) electrode on the left dorsolateral prefrontal cortex and the cathode (negative) electrode on the right dorsolateral prefrontal cortex (F3 and F4 respectively, 10/20 system of electrode placement). In the real/active tDCS condition, stimulation (20 minutes at 2mA) was applied while the participant sat at rest. In the sham condition participants received 30 s of direct current, followed by impedance control with a small current pulse every 550 ms (110 μA over 15 ms) instead of the stimulation current, resulting in an instantaneous current of not more than 2 μA. This method of sham stimulation produced the physical sensations typical of real tDCS and displayed realistic impedance values on the device display. The experimenter was thus blind to the stimulation condition, facilitated by a ‘study’ mode for blinding on the device.

### Image Acquisition

Blood oxygenation level dependent (BOLD) contrast functional images were acquired with echo-planar T2*-weighted (EPI) imaging using a Siemens 3T Magnetom TrioTim syngo MRB17 with a head coil gradient set. Each image was made up of 45 interleaved 3mm thick slices, inter-slice gap, 1mm, field of view 25×25cm; matrix size, 64 × 64; flip angle 87° echo time (TE), 30ms; voxel bandwidth, 2368 Hz/Px; acquisition time (TA), 2.3 s; repetition time (TR), 2710ms. Slice acquisition was interleaved and covered the whole brain with an additional z shim to reduce distortion in the orbitofrontal cortex. Data were acquired in 3 scanning runs of ~5 min each. The first 5 volumes of each run were discarded to allow for T1 equilibration effects.

### FMRI data analysis

FMRI data processing was carried out using FEAT (FMRI Expert Analysis Tool) Version 6.00, part of FSL (FMRIB Software Library, <u>www.fmrib.ax.ac.uk/fsl</u>). Registration to high resolution structural and standard space was carried out using FLIRT(35, 36). Registration from high resolution structural to standard space was then further refined using FNIRT nonlinear registration(37) and motion correction was carried out with MCFLIRT, applying rigid-body transformations. Regressors for each condition yoked to trial onset were convolved with the canonical hemodynamic response function. An intermediate analysis was first carried out, combining three runs into a single dataset for each participant for each testing session (2 per participant). A within-subjects analysis was performed and Z (Gaussianised T/F) statistic images were thresholded using clusters determined by Z>2.3 and a (corrected) cluster significance threshold of p=0.05. Small volume corrected analyses were carried out in bilateral amygdala regions of interest (ROIs). The amygdala ROIs were defined using the Harvard-Oxford Cortical Structural Atlas (using a standard threshold of including all voxels with a greater than 50% probability of lying within the amygdala). The hypothesis-driven contrasts analysed are specified in Table 1:

**Table 1.** Hypothesis-driven contrasts: High level contrasts test for effect of tDCS. Low level contrasts test for effect of task manipulations. tDCS= transcranial direct current stimulation.

| Hypothesis | Analysis | High level contrast | Low level contrast |
| --- | --- | --- | --- |
| Replication of right amygdala response to threat in low attentional load compared to high load. | ROI | Sham only | (Low load (Fear – Neutral)) – High load (Fear – Neutral)) |
| TDCS reduces amygdala response to threat in low attentional load. | ROI | Sham – active | Low load (Fear – Neutral) |
| TDCS increases cortical response to threat in low attentional load. | Whole brain | Active – sham | Low load (Fear – Neutral) |

## Results

Since a previous investigation across a range of trait anxiety levels (24) using this task found that the right amygdala responds selectively on trials with fearful (versus neutral) distractors, when attentional load is low (versus high), we first tested whether this baseline effect was replicated in the present sham condition. As predicted, right amygdala activation occurred selectively on trials with fearful (not neutral) distractors, only when attentional load was low (not high) (load _low-high_ × emotion _fear-neutral_, Z = 3.06, p = .0356, small volume correction (svc)) (Fig. 2-b). This confirmed that our emotional task was sufficiently sensitive to detect the expected presence of amygdala threat signalling in this high anxious sample.

**Figure 2.**
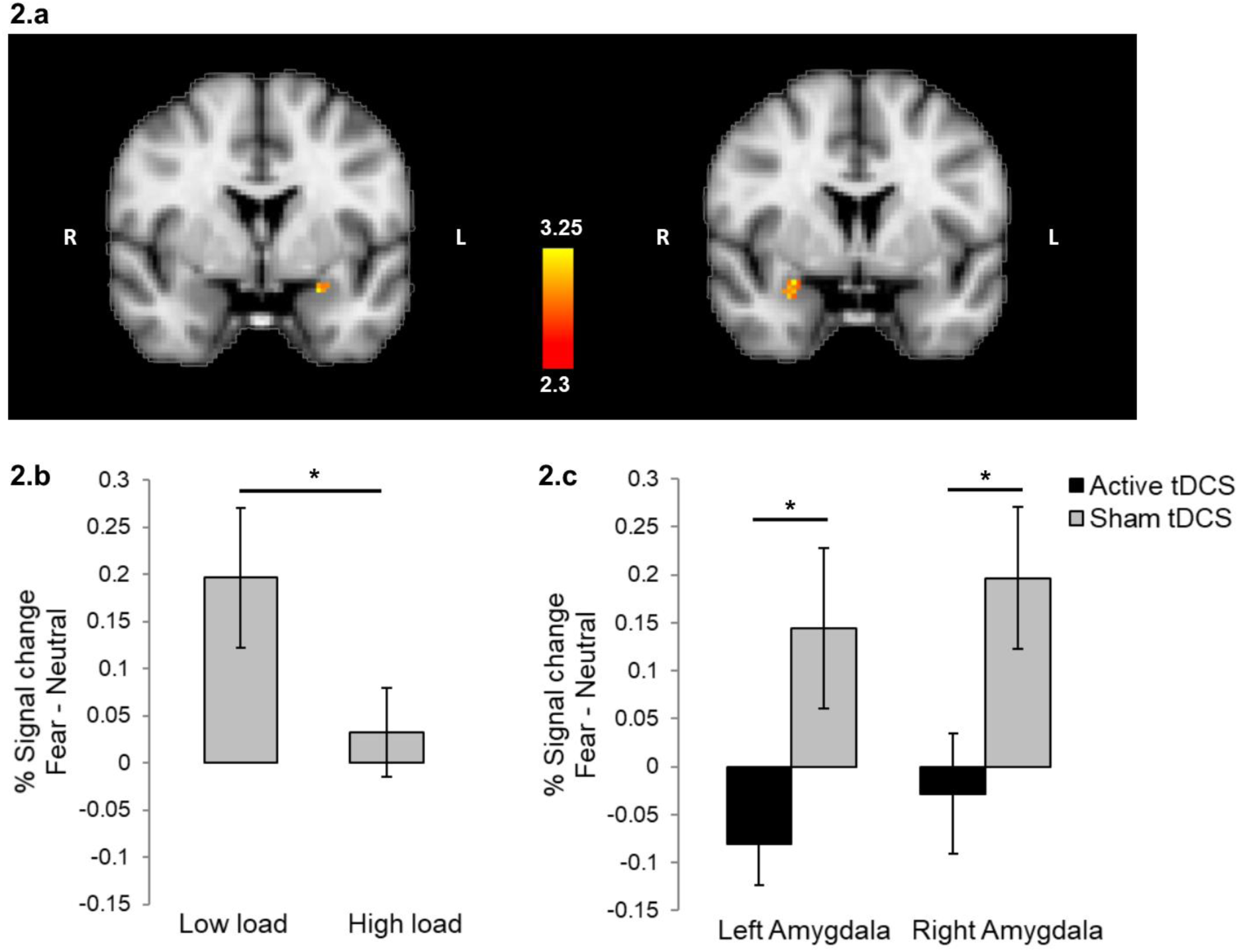
(a) TDCS reduces bilateral amygdala response to fearful faces. Thresholded statistical map depicts clusters in the right and left amygdala (Z>2.3, svc) showing reduced response to fearful versus neutral face distractors under conditions of low attentional load for active compared to sham tDCS (within subjects). Amygdala ROIs defined using the Harvard-Oxford Cortical Structural atlas (thresholded at 50% probability). **(b)** Right amygdala percent signal change extracted and plotted from the significant interaction of fearful face distractors versus neutral face distractors under conditions of low versus high attentional load in the sham (i.e. baseline) condition. **(c)** Right and left amygdala percent signal change data extracted and plotted for the significant cluster shown in (a) for sham versus active tDCS, fearful versus neutral face distractors under conditions of low attentional load. The functional maps are overlaid on an average anatomical image spatially normalised to MNI space. Error bars represent two standard errors of the mean. Asterisks denote statistical significance (* p<.05). N=16

This baseline amygdala response was altered by tDCS. We tested for the predicted effect of tDCS: reduction of amygdala signal for the contrast of fearful-neutral distractors under low load. ROI analysis revealed that in the low load condition only, bilateral prefrontal cortex stimulation significantly *reduced* right amygdala threat response (tDCS _real-sham_ × emotion _fear-neutral_, Z= 3.295, p = .0397, svc), (Fig. 2.a/c.). A similar reduction was observed in left amygdala (Z = 2.816, p = .0401, svc), (Fig. 2.a/c.). Consistent with the absence of a baseline effect of fear in high load trials, tDCS did not modify amygdala activation to fearful versus neutral faces in the high load condition (tDCS _real-sham_ × emotion _fear-neutral,_ (Z<2.3, p>.05)). Rather, tDCS reduced amygdala response to fearful versus neutral distractors selectively in the low load condition.

Whole brain voxelwise analysis revealed that, contrary to the amygdala effect, stimulation significantly *increased* activation in cortical regions associated with attentional processing (tDCS _real-sham_ × emotion _fear-neutral_: right supramarginal gyrus, right superior temporal sulcus: Z = 3.471, p = .0005, whole brain corrected; Left superior frontal sulcus, precentral gyrus, frontal eye fields: Z = 3.74, p = .0066, whole brain corrected), (Fig. 3.a/b.). See table 2 for results summary. Consistent with the absence of a baseline effect of fear, and no effect of tDCS on high load trials, tDCS did not modify whole brain activation to fearful versus neutral faces in the high load condition (tDCS _real-sham_ × emotion _fear-neutral_, (Z<2.3, p>.05)). Rather, tDCS increased cortical response to fearful versus neutral distractors in the low load condition.

**Table 2.**
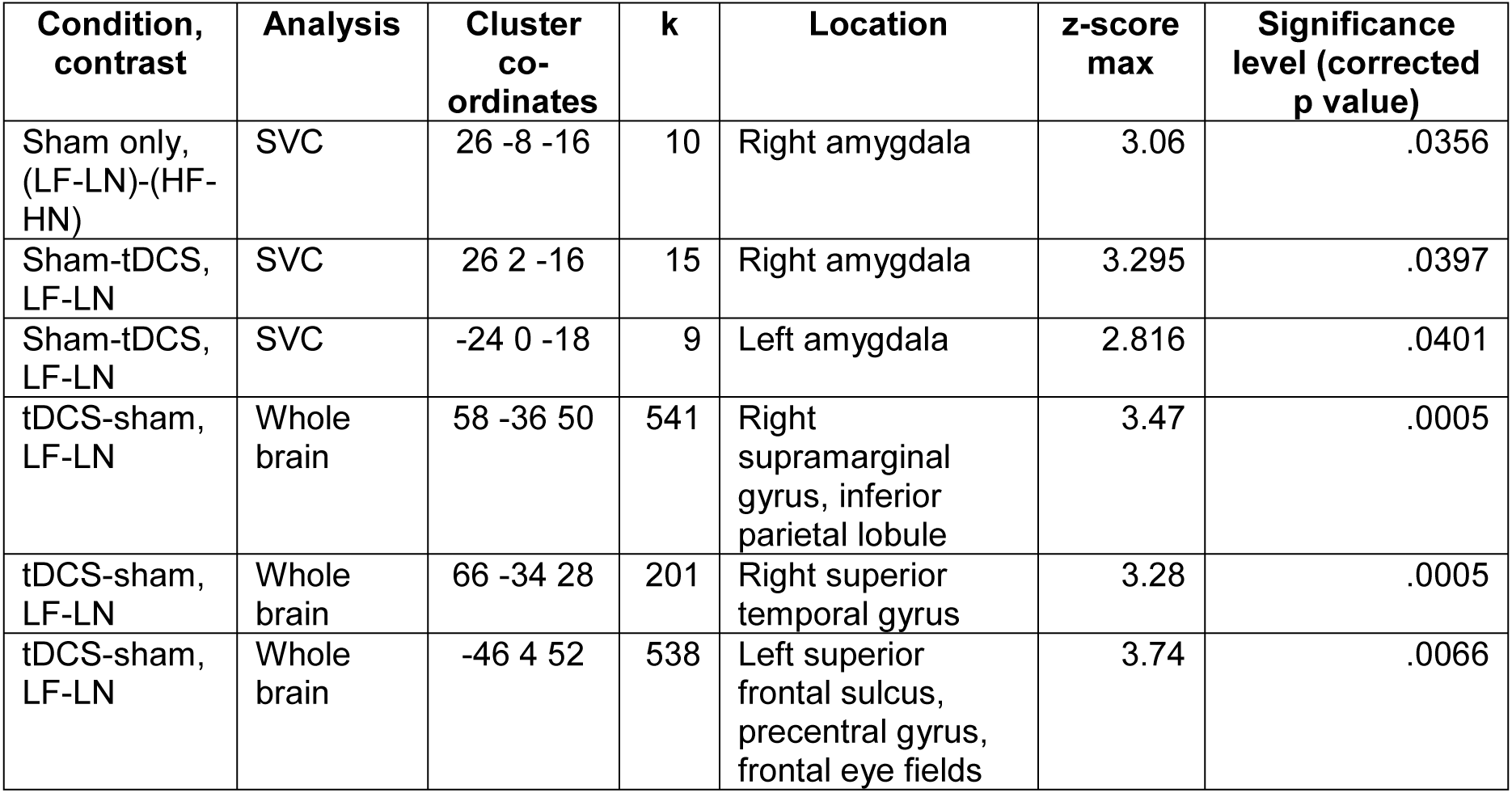
Key findings summary: Hypothesis refers to Table 1 above. LF= Low attentional load, fearful face, LN= Low attentional load, neutral face, HF= High attentional load, fearful face, HN= High attentional load, neutral face. SVC= Small volume correction.

| Condition, contrast | Analysis | Cluster co-ordinates | k | Location | z-score max | Significance level (corrected p value) |
| --- | --- | --- | --- | --- | --- | --- |
| Sham only, (LF-LN)-(HF-HN) | SVC | 26 -8 -16 | 10 | Right amygdala | 3.06 | .0356 |
| Sham-tDCS, LF-LN | SVC | 26 2 -16 | 15 | Right amygdala | 3.295 | .0397 |
| Sham-tDCS, LF-LN | SVC | -24 0 -18 | 9 | Left amygdala | 2.816 | .0401 |
| tDCS-sham, LF-LN | Whole brain | 58 -36 50 | 541 | Right supramarginal gyrus, inferior parietal lobule | 3.47 | .0005 |
| tDCS-sham, LF-LN | Whole brain | 66 -34 28 | 201 | Right superior temporal gyrus | 3.28 | .0005 |
| tDCS-sham, LF-LN | Whole brain | -46 4 52 | 538 | Left superior frontal sulcus, precentral gyrus, frontal eye fields | 3.74 | .0066 |

**Figure 3.**
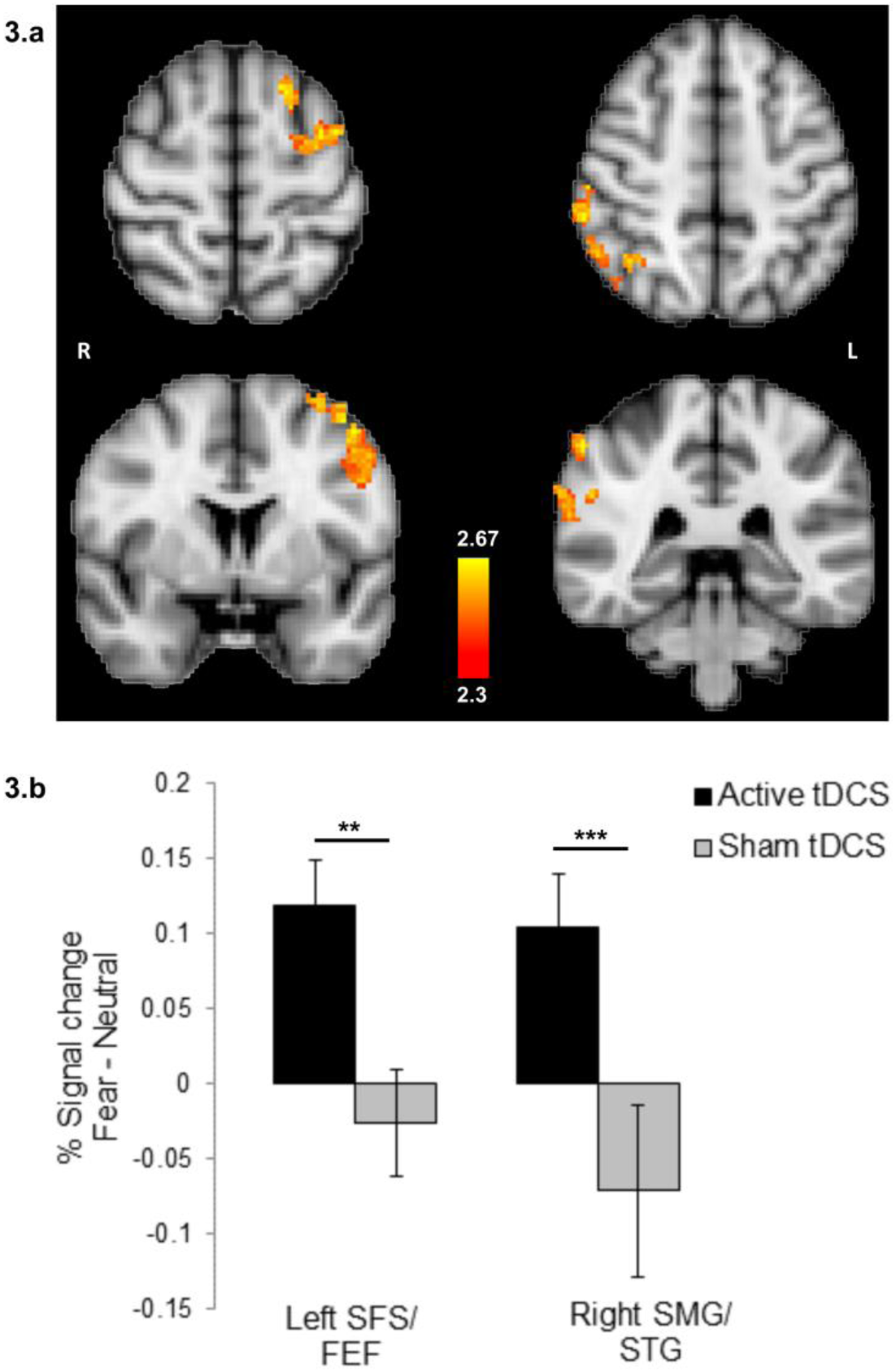
(a) TDCS increases cortical activation in response to fearful faces. Thresholded statistical maps (Z>2.3, whole brain corrected) depict significant clusters in cortical regions associated with attentional processing (left superior frontal sulcus, precentral gyrus, frontal eye fields and right supramarginal gyrus, superior temporal sulcus), for active tDCS compared to sham for the contrast of fearful-neutral face distractors under low attentional load **(b)** Percent signal change extracted and plotted for the significant clusters shown in **(a)** for active versus sham tDCS, for the contrast of fearful-neutral face distractors under low attentional load. The functional maps are overlaid on an average anatomical image spatially normalised to MNI space. Error bars represent two standard errors of the mean. Asterisks denote statistical significance (** p<.01, *** p<.001). N=16

Next, we tested whether prefrontal stimulation would also improve anxious participants’ task performance under threatening conditions. To assess this, we contrasted task accuracy following real and sham tDCS. The results of higher level contrasts were not significant (e.g. tDCS × load × emotion, all p>.05). However, in the key condition (low load, fearful trials) the contrast of real – sham tDCS indicated improved accuracy after stimulation (mean difference: 2.2 (9.1%), SD = 4.5; t(15)=1.936, p=.036,1-tail, dz = 0.49).

## Discussion

As predicted, in this high trait anxious female sample, prefrontal stimulation remediated fronto-limbic governance of attentional control under threat. Functionally, stimulation simultaneously reduced amygdala and increased cortical activation to fearful face distractors. Behaviourally, this was accompanied by reduced influence of distractors on task accuracy – i.e. reduced attentional capture by threat. The demonstration that prefrontal cortex stimulation can abolish amygdala threat reactivity, and simultaneously increase activity in cortical regions associated with attentional control(38), provides the first experimental evidence for a direct causal inhibitory role of prefrontal cortex on amygdala threat response in human trait anxiety. These findings suggest a mechanism of action that may contribute to treatment effects of tDCS previously observed in clinical trials of affective disorders.

The amygdala is one of the key brain regions implicated in the pathophysiology of depression and anxiety disorders. Pre-clinical research in the 1980s showed that conditioned fear was mediated by the transmission of information to the amygdala, and that control of fear reactions was mediated by output projections from the amygdala to the behavioural, autonomic, and endocrine response control systems in the brainstem(39). Inactivation of the amygdala during fear conditioning, through focal amygdala infusions, has been shown to prevent the acquisition and expression of fear conditioning(40). Depressed patients exhibit hyperactive amygdala responses to emotional information (for a review, see (9)), as do patients with anxiety disorders(41, 42). Treatment with selective serotonin reuptake inhibitors (SSRIs) has been shown to attenuate amygdala response to fearful faces in healthy controls (43) and in depressed patients(44), both before reported treatment response(45) and after (46). The latter studies indicate that a reduction in amygdala hyperactivity may be an important part of the mechanism of action of SSRIs in improving mood.

Animal studies suggest that, at least in the case of conditioned fear responses, the amygdala response to threat can be extinguished by top down inhibition from the medial prefrontal cortex(4). In humans, brain imaging studies of anxiety have revealed hypo-activation of the lateral prefrontal cortex(47), particularly in the context of fearful distractors(48), which is thought to reflect deficient attentional control. However, fMRI is a correlational technique and there has been no causal test of the relationship between frontal and amygdala activity in humans, especially as pertaining to attentional capture by threat. The present study demonstrates that modulating excitability in the prefrontal cortex can significantly reduce the amygdala response to threat-related distractors.

Our prior behavioural investigation with prefrontal tDCS indicates that it has the potential to reduce vigilance to threat in a paradigm reported to predict the clinical response to anxiolytic treatment(23). Here we investigated a possible neural mechanism that may mediate this effect: increased suppression of amygdala threat responsivity through improved regulation from a top-down attentional control network. As well as amygdala hyperactivity, depressed(9) and anxious(7) patients have decreased frontal activation in response to cognitive tasks(8). Previous work(24) has found that high anxious participants show increased amygdala and decreased frontal activation to fearful distractor faces under conditions of low attentional load, compared to low anxious participants. Our high anxious sample showed a similar profile of amygdala response to fearful versus neutral face distractors under low attentional load in the sham stimulation condition (Fig. 1). As hypothesised, we found that prefrontal cortex tDCS reduced this activation, such that after stimulation our high anxious group more closely resembled the low anxious participants in the prior study, with increased cortical and reduced amygdala response to threat distractors under low load. Furthermore, this was accompanied behaviourally by reduced attentional capture by threat distractors under low load, reflected in increased accuracy levels following tDCS.

Another form of brain stimulation, transcranial magnetic stimulation (TMS), of prefrontal cortex is an FDA-approved treatment for depression(49). By contrast, the evidence base for clinical efficacy of tDCS is still in development. If efficacy is established, then tDCS offers several potential advantages over TMS, including being better tolerated, and cheaper/simpler to administer, while the development of home use devices broadens both potential patient uptake and clinical research. One clinical trial in a sample of 120 depressed patients compared repeated daily treatment with DLPFC tDCS for three weeks versus treatment with the antidepressant drug sertraline (50 mg/day)(50). The results suggested that the combined effects of tDCS and sertraline relieved depressive symptoms more quickly and effectively than either treatment alone. In addition, tDCS showed a similar level of efficacy to sertraline, but only tDCS (and not sertraline) was superior to placebo in this trial. These results highlight the potential value of tDCS in the treatment of mood disorders. However, as with many antidepressant drug treatments, the mechanism of action of tDCS is unclear. The present proof-of-concept study used a single-session tDCS intervention, and describes a neurocognitive mechanism of action of prefrontal tDCS in trait anxiety that should be investigated in future therapeutic trial designs: clinical efficacy as a function of reducing hyperactive amygdala-dependent vigilance to threat.

In the present study, the task was conducted minutes after the end of stimulation. Therefore, it is conceivable that, in some individuals, the physiological after-effect of tDCS may have already decayed significantly during task performance. The duration of after-effects of prefrontal tDCS has not been studied systematically. However, in motor cortex, the same stimulation protocol used here (2mA, 20 minutes) had physiological after-effects lasting for at least 90 minutes after stimulation offset(51). In our study, the attentional control task was carried out over 40 minutes, starting ~7 min after stimulation offset. If one can presume to generalize from motor to prefrontal cortex, then testing was conducted within the likely time window of tDCS physiological after-effect. The purpose of the present study was to investigate acute effects of single-session tDCS, by testing for an induced change in amygdala response to threat. Future work is required to determine whether these effects are extended over time when repeated tDCS interventions are used, and to determine if these acute changes in neurobehavioural markers of threat vigilance are predictive of treatment response. To effectively build upon this initial investigation of the neural mechanisms of tDCS, future imaging work should examine changes in functional connectivity between the dorsal attention network, the amygdala and other regions implicated in treatment response. To increase homogeneity of the sample and because of higher prevalence of high trait anxiety and anxiety disorders in females(52), only females were recruited for this study. Future work is required to investigate sex differences and generalizability to the male population.

A number of recent studies have indicated that repeated (10 sessions over 2 weeks) administration of prefrontal tDCS may be an effective treatment for depression(50, 53). The current results from a single session in anxious participants reveal an effect of prefrontal tDCS on a neural biomarker relevant to clinical depression and anxiety. Taken together with our prior behavioural findings with the same stimulation protocol(23), this indicates a potential neurocognitive mechanism (reduced fear vigilance, facilitated by increased cortical and reduced amygdala activation) that may partially mediate the reported clinical efficacy of prefrontal tDCS in the literature. This neural biomarker may have potential to test and benchmark novel stimulation protocols in the development phase, to optimize treatment efficacy for depression and anxiety. Confirming preclinical animal studies, and correlational evidence from human neuroimaging, the present study offers the first causal evidence in humans of the relationship between prefrontal cortex and the amygdala in regulating threat processing in trait anxiety.

## Acknowledgements

This research was funded by an MRC studentship awarded to Maria Ironside. Sonia J Bishop is supported by ERC Starting Grant No 260932 and NIH grant R01MH091848. JO’S was supported by the National Institute of Health Research Oxford Biomedical Research Centre (NIHR BRC). The views expressed are those of the authors and not necessarily those of the NHS, the NIHR or the Department of Health.

